# The determinants of alpine butterfly richness and composition vary according to the ecological traits of species

**DOI:** 10.1101/002147

**Authors:** Vincent Sonnay, Loïc Pellissier, Jean-Nicolas Pradervand, Luigi Maiorano, Anne Dubuis, Mary S. Wisz, Antoine Guisan

## Abstract

Predicting spatial patterns of species diversity and composition using suitable environmental predictors is an essential element in conservation planning. Although species have distinct relationships to environmental conditions, some similarities may exist among species that share functional characteristics or traits. We investigated the relationship between species richness, composition and abiotic and biotic environment in different groups of butterflies that share ecological characteristics. We inventoried butterfly species richness in 192 sites and classified all inventoried species in three traits categories: the caterpillars diet breadth, the habitat requirements and the dispersal ability of the adults. We studied how environment, including influence butterfly species richness and composition within each trait category. Across four modelling approaches, the relative influence of environmental variables on butterfly species richness differed for specialists and generalists. Climatic variables were the main determinants of butterfly species richness and composition for generalists, whereas habitat diversity, and plant richness were also important for specialists. Prediction accuracy was lower for specialists than for generalists. Although climate variables represent the strongest drivers affecting butterfly species richness and composition for generalists, plant richness and habitat diversity are at least as important for specialist butterfly species. As specialist butterflies are among those species particularly threatened by global changes, devising accurate predictors to model specialist species richness is extremely important. However, our results indicate that this task will be challenging because more complex predictors are required.

## Introduction

Species distribution and abundance patterns are highly sensitive to changes in climate (Thuiller *et al*. 2005) and habitat degradation (Donald, Green & Heath 2001; Hewison *et al*. 2001). Species are expected to respond distinctively to these changes, and recent investigations demonstrate that important differences between taxa, related primarily to different ecological traits, can influence the responses of species to environmental changes (Henle *et al*. 2004; Syphard & Franklin 2009). For example, Williams et al. (2010) observed that social bee species were more affected by isolation from natural habitat and pesticides than solitary species, whereas habitat specialists are more vulnerable to habitat degradation than habitat generalists (plants: Fischer and Stocklin 1997, butterflies: Warren et al. 2001, mammals: Fisher et al. 2003, carabid beetle: Kotze and O’Hara 2003, birds: Julliard et al. 2004). As a consequence, the ability to protect biodiversity requires a good understanding of how species ecology relates to the drivers of biodiversity.

Spatial variation of biodiversity correlate with numerous environmental factors (Gaston 2000). Examples of drivers include temperature (Stevens 1989; Gaston 1996), ambient energy (Wright 1983; Currie 1991; Hawkins *et al*. 2003), habitat heterogeneity (Shmida & Wilson 1985; Kerr & Packer 1997), and land cover (Nogues-Bravo & Martinez-Rica 2004). These relationship are complex, as the role of these drivers may change across spatial scales (e.g., Rahbek and Graves 2000, Lortie et *al.* 2004), but also according to the species ecological specialisation For instance, Ribera et al. (2001) established that ground beetles with low dispersal ability are more sensitive to land disturbances than mobile species. Thus, to obtain a better understanding of the drivers of community composition and diversity, studying the distribution of taxa along environmental gradients requires consideration of the species ecology.

Butterflies exhibit a diverse range of diet, habitat requirements and dispersal abilities (Lopez-Villalta 2010). As is the case in other groups of herbivorous invertebrates, caterpillars have developed a large range of trophic specialisation, ranging from strictly monophagous to polyphagous species (Ehrlich & Raven 1964, 1967). The habitat occupied by the adult is related to the larval diet requirements, at least for butterflies with limited dispersal abilities. Some species specialize on a restricted number of habitats and exhibit only limited dispersal within these habitats (Warren *et al*. 2001). In contrast, more generalist species frequently display higher dispersal ability and are able to move from one habitat type to another (Warren *et al*. 2001). Consequently, variation in ecological traits is expected to affect butterfly species richness and composition in communities by influencing how butterfly species respond to environmental conditions.

Many studies have demonstrated strong positive correlations between butterfly species richness, temperature and solar radiation, along with a negative correlation with rainfall (Turner, Gatehouse & Corey 1987; Pollard 1988; Roy *et al*. 2001; Luoto *et al*. 2006; Illan, Gutierrez & Wilson 2010). These findings support hypotheses that link net available energy with the metabolic needs of ectothermic species. Because butterfly larvae frequently depend on particular host plant species, plant species richness and composition have also been identified as important predictors of butterfly species richness and composition, depending on the scale considered, (Kerr, Southwood & Cihlar 2001; Hawkins & Porter 2003a; Menendez *et al*. 2007). An increased range of resources can potentially sustain higher butterfly diversity (Erhardt 1985), and decreasing plant diversity has been shown to correlate negatively with butterfly diversity (Illan *et al*. 2010; Stefanescu, Carnicer & Penuelas 2011). Soil nitrogen content or soil acidity, are also known to affect butterfly species richness and composition indirectly by modifying the vegetation structure (Vinton & Burke 1995; Roem & Berendse 2000). Ecological traits can affect the way those environmental conditions influence butterfly diversity. In Britain, lowland climate has a strong effect on habitat-generalist butterflies, whereas host-plant richness and habitat heterogeneity are more important for specialist species (Menendez *et al*. 2007). In different geographical regions of Finland, Ekroos et al. (2010) showed that intensive land use practices impacting plant communities caused homogenisation of butterfly communities and especially a decrease in specialist species.

The aim of this study was to establish how ecological traits influence the response of butterfly species richness and composition to environmental conditions. This analysis can suggest how species with different characteristics may cope with global change, but it can also indicate whether the distribution of generalist and specialist species can be modelled with similar accuracy for use as a conservation tool. Accordingly, we investigated the relationship between butterfly species richness, composition and climatic, landscape and vegetation variables in the western Swiss Alps as a function of species diet, dispersal ability and habitat requirements.

## Methods

### Study area

The study area is located in the Western Swiss Alps (Figure 1), with calcareous soils, a temperate climate (Bouët 1985), and elevations ranging from 1,000 to 3,210 m a.s.l. We selected sampling plots following a balanced stratified sampling design based on altitude, slope and aspect (Hirzel & Guisan 2002) and considering only open, non-forested areas. A two-year sampling effort (2009 and 2010) allowed us to visit a total of 192 plots (50 m x 50 m each). Each plot was visited every three weeks between June 1 and September 15. Following Pollard et al. (1993), we sampled only during good weather conditions (low wind, sunny and high temperature) and between 10 am and 5 pm, when butterflies are most active. In each plot, we used a net to collect all butterflies (Papilionoidea, Hesperiidae, and Zygenidae) present during a 45-minute sampling period. All butterflies were identified directly in the field to the species level. Moreover, we conducted an exhaustive vegetation inventory covering 4 m^2^ in the center of each plot.

**Figure 1:**
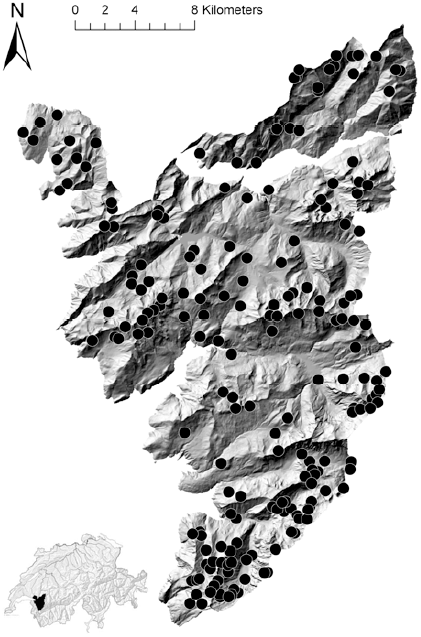
Study area in the Western Swiss Alps. The black dots represent the locations of the butterfly plots sampled.

### Species trait data

We classified all butterfly species observed in the field into different classes. The classes were defined by the ecological traits of the species, with a particular focus on diet, habitat requirements and dispersal ability. We collected data on diet and habitat requirements from LSPN (1987). Based on the diet of the caterpillars, we classified all species as either diet specialists (those whose caterpillars use from one to three host plant species) or diet generalists (those that use more than three host plant species). The limit defined at three host plant species was selected in order to create two categories of more or less equal sample size. To define the host-plant specialisation, we preferred to use the number of host-plant species rather than more frequently used index based on the plant taxonomy (e.g. number of plant genera). The number of plant species is a better indication of the possibility for a butterfly to find a suitable oviposition site in the landscape. We obtained 61 diet specialist species and 70 diet generalist species. We defined three classes based on the habitat used by the adults (see Table S1 in supporting information): habitat specialist species, those restricted to one or two different habitats (n=58); habitat intermediate species, those found in up to four habitats (n=42); and habitat generalist species, those found in more than four habitats (n=31). The arbitrary limits were also selected in order to obtain comparable sample size in each category. We also defined two classes based on dispersal ability (Bink 1992): low dispersal ability (n=84) and vagile species (n=47).

### Environmental predictors

To study the pattern of butterfly species richness in the study area, we selected a set of seven environmental variables specifically related to butterfly ecology. We considered two topoclimatic variables (temperature and solar radiation), two landscape variables (the proportion of forests and the Shannon habitat diversity index), two pedologic variables (soil nitrogen content and soil acidity), and plant species richness.

Temperature strongly affect butterfly distribution (Luoto *et al*. 2006), essentially because it is tightly linked to the ability of the pupae to emerge as an imago and to the physiological tolerance to freezing. Because caterpillar development can be represented by a degree-day model (van Asch & Visser 2007), we calculated degree-days (DDEG), based on monthly average temperatures following Zimmermann and Kienast (1999), to indicate the effect of temperature. Solar radiation (SRAD) has been viewed as one of the main drivers of butterfly richness (Turner *et al*. 1987), and we calculated this variable as in Kumar et al. (1997). Both DDEG and SRAD were calculated at a resolution of 100 m. The two landscape variables, calculated from the land cover data provided by the Swiss Federal Office of Statistics, were considered because of the positive relationship that exists between spatial environmental heterogeneity and the species richness of arthropods (Hendrickx *et al*. 2007). We calculated the proportion of forest (PROPFOR) and the habitat diversity (HABDI) at a resolution of 100 m using the software FRAGSTAT (McGarigal & Marks 2000), with a mobile window with a 500 m radius. The two pedologic variables were derived from Landolt ecological values (Lauber & Wagner 2007): an index of soil nitrogen content (N) and an index of soil acidity (PH) similar to that developed by Ellenberg (1988) for central Europe. For each sampled plot, both indexes were calculated as the mean nutritive substance value and the mean reaction value of the plant species inventoried; this method that has been proven reliable in other studies (Scherrer & Korner 2011).

### Species richness

We built richness models for all butterfly species together as well as for each category separately. Given that different approaches to modelling species distributions generate significant variability in the results (Segurado & Araujo 2004) and produce potentially different answers (Marmion *et al*. 2009), we considered four different statistical methods to assess whether the different approaches were consistent: generalised linear models (GLMs) (McCullagh & Nelder 1989), generalised additive models (GAMs) (Hastie & Tibshirani 1990), generalised boosted models (GBMs) (Ridgeway 1999) and random forests (RFs) (Breiman 2001). We used a Poisson distribution. We allowed a maximum of 3000 trees, and we used a maximum of 3000 trees for the construction of the RFs. To test the prediction accuracy of the models, we applied a repeated (100 times) data-splitting procedure (Dubuis *et al*. 2011) to each model. For each run, we split the original dataset to obtain a 70%-30% partition. The 70% partition was used to fit the models. The remaining 30% was used for independent evaluation of the models. Then, for each split-sample repetition and for each model, we calculated a Spearman correlation coefficient between the observed and predicted species richness using the evaluation data set. This correlation measured the predictive power of the model. We also evaluated the reliability of the GLMs and the GAMs by calculating the adjusted deviance (i.e., the explanatory power of the model).

Breiman (2002) suggested that the reduction in mean squared error (MSE) should be used to investigate the importance of variables in tree-based modelling techniques involving permutations of variables. This approach, termed permutation-based MSE reduction, is now accepted as the state-of-the-art method (Grömping 2009). Permutation-based MSE reduction has the additional advantage that it can be applied to standard regression techniques to quantify the importance of variables for comparative purposes (Thuiller *et al*. 2009). To quantify the contribution of each variable to the fitted model, we applied a similar permutation procedure to each model using 100 iterations as described in Thuiller et al. (2009). To assess the importance of a given variable, this procedure uses a Spearman correlation between the standard predictions and the predictions obtained from random permutations of the variable under investigation. This approach is based on the principle that the model need not necessarily contain all the variables considered, provided that the prediction remains effective (Grömping 2009). Thus, a slight reduction in the correlation value suggests that the variable in question is of little importance for the predictive power of the model.

### Communities composition

We run canonical correspondence analyses (CCA) for each trait category separately. This analysis relates the species composition with the seven previously mentioned environmental factors (Ter Braak 1986, 1987). We use the package ade4 implemented in the R environment. We calculated the variance explained by each axis separately.

## Results

During the two years of sampling, we observed a total of 131 butterfly species belonging to 59 different genera. Forty-seven species were found on the richest plot, whereas none were found on the poorest plot. A total of 562 plant species were identified in the vegetation plots. The richest plot contained 69 plant species, and the poorest plot contained two plant species.

Topoclimatic predictors had the greatest overall ability to explain the species richness patterns if all species were considered together (Figures 2). Degree-days and solar radiation had the greatest explanatory power for diet and habitat generalist species and for species with high dispersal ability. The influence of the environmental predictors on species richness did not differ between habitat specialists and habitat-intermediate species. In contrast, plant species richness and habitat diversity had greater explanatory power than degree-days and solar radiation for habitat specialists, for diet specialist species and species with low dispersal ability. To a lesser extent, the soil acidity also contributed to the richness of habitat specialist species. However, it did not affect the richness of generalist species. In all cases, the proportion of forest and the soil nitrogen had only a very weak explanatory power for butterfly species richness.

**Figure 2:**
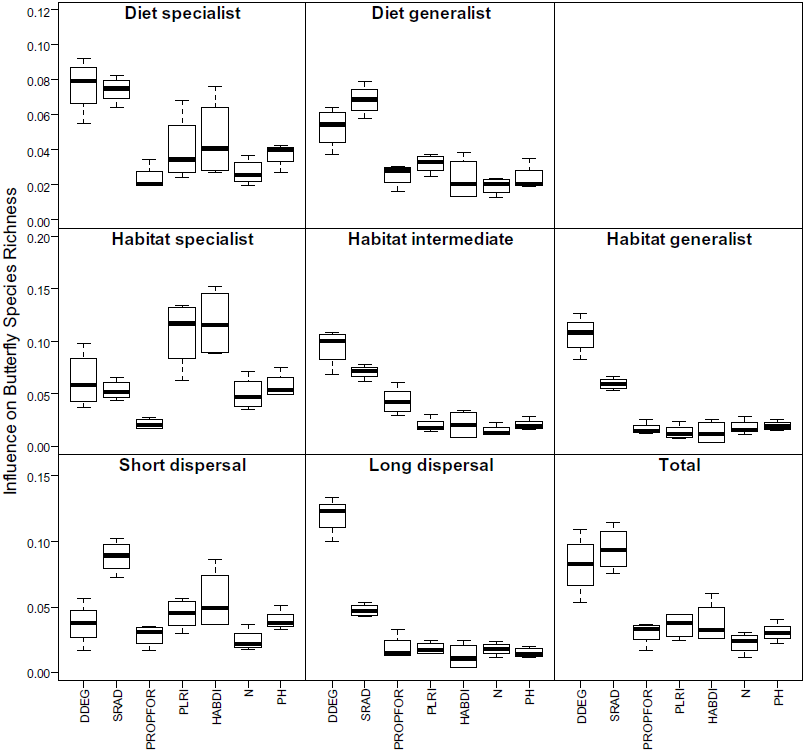
Influence of different environmental predictors – degree-days (DDEG), solar radiation (SRAD), proportion of forest (PROPFOR), plant species richness (PLRI), habitat diversity (HABDI), nitrogen soil content (N) and soil acidity (PH) – on butterfly species richness according to diverse ecological trait classifications.

The results of the analyses of predictive power indicated that better predictions were obtained for species with high dispersal ability (0.737±0.008) than low (0.657±0.022) or intermediate habitat requirements (0.689 ± 0.011) than for species with low dispersal ability (0.499±0.014) or high habitat requirements (0.463±0.053). No large differences in the predictive power of the models were found between diet specialist and diet generalist species. Each of the four modelling approaches yielded very similar predictive power and the relative importance of each predictor (see Table S2 in supporting information).

Results provided by the CCA supported those obtained on species richness (Figure 3, and see Table S3 and S4 in supporting information). The variance explained by the two first axis of the CCA is greater for generalist species communities (whether it concerns habitat – 75% or diet 78%) than for specialist species communities (respectively 64% and 53%). Moreover, this difference is almost exclusively supported by the first axis which represents a proportionally greater part of the explained variance in the case of generalist species communities than in the case of specialist species communities. The first axis mainly corresponds to climatic variable. No major difference was found between species with long and short dispersal ability.

**Figure 3:**
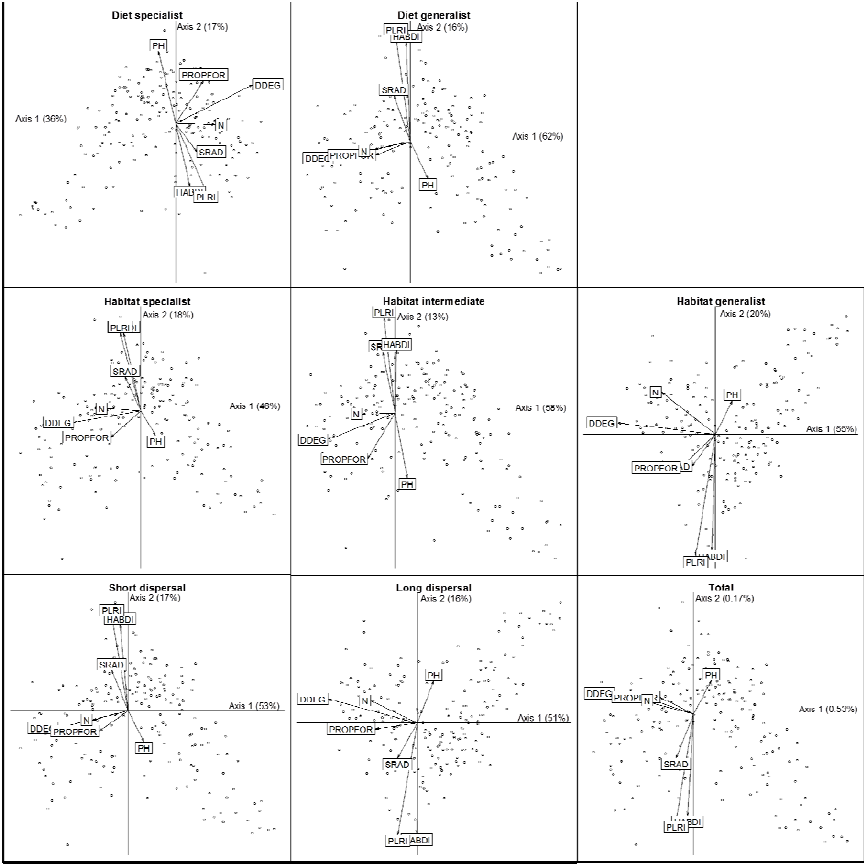
Canonical correspondence analysis ordination diagram representing the species composition of the 192 sampling plots (dots) along with the different environmental predictors (arrows). Only the two first axes are presented.

## Discussion

Biodiversity patterns and the main drivers affecting species richness are increasingly well documented in fauna and flora worldwide (Gaston 2000). Here, we show how species displaying more or less specialized ecology are driven by environmental predictors that differ from those that drive the ecology of more generalist species. Degree-days and solar radiation are the most important predictors of butterfly richness and composition for ecologically generalist species. For specialist species, plant species richness and habitat diversity, have a much higher importance. Because capturing the environment of specialist species is more difficult and require more complex predictors (e.g. biotic variables), it appears that predictions of species richness are less reliable for specialist species than for generalist species.

Arthropod diversity and plant richness are expected to be strongly correlated (Siemann 1998; Siemann *et al*. 1998). However, Stefanescu et al. found that the number of plant vegetation type was not related to butterfly richness (2004) or that it was only a poor predictor of butterfly richness (2011). In contrast, our results provide strong evidence that plant species richness affect the richness and composition of specialist butterflies. Similarly, Illán et al. (2010) found, in a study conducted in Spain, that land cover had only a marginal effect on butterfly diversity, contrasting with our finding that the habitat diversity index influences the diversity of specialist butterfly species. These contrasting observations may indicate that very proximal predictors are necessary to model specialist butterfly species richness. However, creating proximal predictors requires a large amount of data and ecological knowledge limiting the prediction accuracy. Indeed, we found that prediction accuracy was lower for habitat specialists and species with low dispersal ability than for habitat generalists and species with higher dispersal ability. Identifying and developing variables useful for modeling specialist species is worthwhile because those species are often the most threatened (Belgium: Polus *et al*. 2006, UK: González-Megías *et al*. 2008, Finland: Poyry *et al*. 2009, Spain: Stefanescu *et al*. 2010), and reliable models could help inform conservation efforts.

A growing body of evidence indicates that modeling species distribution and community composition requires both abiotic and biotic variables (e.g. Pellissier *et al*. 2010; Wisz *et al*. in press, Pellissier *et al*. in press). Here, because caterpillars can be specialized on a low number of host-plant species occurring in specific habitats, the distribution of those species can be largely constrained. The association that we found between habitat diversity, plant species richness and the richness of specialist species of butterflies emphasizes the importance of biotic interactions or more specifically trophic interactions to predict communities (Pellissier et *al.* in press).

Overall, the climate remains the main factor affecting butterfly diversity and composition in alpine landscape. In mountainous regions, climate may impact herbivorous insects in two distinct ways. It constrains species richness directly through the physiological requirements of species (e.g., butterfly larvae cannot grow below a given temperature threshold) and indirectly by affecting resource availability for instance by affecting host-plant availability. We showed that degree-days and solar radiation are both important predictors of butterfly richness for both generalist and specialist species. This clear result is consistent with previous findings that climatic predictors linked with the input of solar energy are the main determinants of the diversity of many groups of organisms (Currie 1991), including butterflies (Turner *et al*. 1987; Hawkins & Porter 2003b; Luoto *et al*. 2006).

In contrast to the findings in Stefanescu et al. (2004, 2011), we found evidence that acidity influences the richness of specialist butterfly species. This finding may result from changes in the composition of the plant community with soil acidity (Partel *et al*. 2004). In particular, soil acidification results in a decrease of the cover of host plant families, such as Fabaceae or Poaceae, that are highly favorable to a large number of butterfly larvae. However, despite its influence on plant community composition, the soil content of nitrogen appeared to be a poor predictor of butterfly richness. Similarly, the proportion of forest was not important in the models. Forest edges constitute a full-fledged environment that is highly favorable for some butterfly species. A forest-edge effect affects butterfly species richness positively and decreases as the distance to the forest increases(Ohwaki *et al*. 2007, Bossart and Opuni-Frimpong 2009, Marini *et al*. 2009). However, no similar effect was observed in our study. In all cases, the proportion of forest appeared to be a poor predictor of trends in butterfly species richness.

We emphasized that the four species distribution models that we used are consistent with one another and yield homogeneous patterns. However, variability between those approaches remains. This variability is almost surely the consequence of mathematical differences among the models. GLMs and GAMs are based on polynomial regressions, whereas GBMs and RFs are based on decision trees. Several recent studies emphasize that uncertainties in statistical approaches should be considered in predictive modelling (Elith, Burgman & Regan 2002; Barry & Elith 2006; Heikkinen *et al*. 2006). Thus, our results are congruent with the recommendations of Marmion et al. (2009) who argue that a multi-method approach should be used for species distribution modelling in order to check the consistency of predictions and of findings regarding the importance of different variables.

In conclusion, we showed that climate variables appear to be the strongest drivers affecting trends in generalist butterfly species richness. However, our findings also indicated that additional variables, such as plant richness and habitat diversity, are at least as important as climate variables for predicting the patterns of species richness of specialist butterflies. The influence of plant richness and habitat diversity demonstrates the importance of incorporating proximal predictor to explain the distribution, richness and composition of specialist species. The species most endangered by global change typically share particular life-history traits (Barbaro & van Halder 2009) and are primarily specialists (Poyry *et al*. 2009). However, our results indicate that these are the species for which it is most challenging to create suitable predictors and predict distributions. Our results therefore highlight the critical need to develop more proximal predictors of specialist species distribution and richness.

**Table 1:** Predictive power (i.e., adjusted deviance) and explanatory power (i.e., data-splitting procedure) values for four different species richness models according to the diverse ecological traits investigated.

| Ecological trait classes |  | Statistical method | GLM | GAM | GBM | RF | Average |
| --- | --- | --- | --- | --- | --- | --- | --- |
| Diet | specialist | Ajusted deviance | 0.452 | 0.515 |  |  |  |
| | | Data splitting procedure | 0.613 | 0.565 | 0.607 | 0.630 | $0.604 \pm 0.028$ |
|  | generalist | Ajusted deviance | 0.426 | 0.477 |  |  |  |
| | | Data splitting procedure | 0.571 | 0.570 | 0.564 | 0.608 | $0.578 \pm 0.020$ |
| Dispersal | short | Ajusted deviance | 0.359 | 0.414 |  |  |  |
| | | Data splitting procedure | 0.503 | 0.498 | 0.481 | 0.514 | $0.499 \pm 0.014$ |
|  | long | Ajusted deviance | 0.597 | 0.638 |  |  |  |
| | | Data splitting procedure | 0.731 | 0.739 | 0.730 | 0.748 | $0.737 \pm 0.008$ |
| Habitat | specialist | Ajusted deviance | 0.278 | 0.347 |  |  |  |
| | | Data splitting procedure | 0.440 | 0.422 | 0.450 | 0.540 | $0.463 \pm 0.053$ |
|  | intermediate | Ajusted deviance | 0.531 | 0.564 |  |  |  |
| | | Data splitting procedure | 0.697 | 0.698 | 0.674 | 0.690 | $0.689 \pm 0.011$ |
|  | generalist | Ajusted deviance | 0.512 | 0.570 |  |  |  |
| | | Data splitting procedure | 0.638 | 0.649 | 0.653 | 0.689 | $0.657 \pm 0.022$ |
| Total |  | Ajusted deviance | 0.474 | 0.523 |  |  |  |
| | | Data splitting procedure | 0.607 | 0.414 | 0.588 | 0.617 | $0.557 \pm 0.096$ |

## Acknowledgements

We thank all those who helped with field data collection. This study was supported by the European Commission (ECOCHANGE project, contract no. FP6 2006 GOCE 036866) and NSF grant no. 31003A-125145 (BIOASSEMBLE project).

## Supplementary materials

**Table S1:** List of the different classes to calculate the species habitat requirements.

| Environment/habitat | Definition |
| --- | --- |
| Nitrogen free grasslands/pastures | Grasslands without regular addition of fertilizers |
| Fertilized grasslands/pastures | Grasslands with regular addition of fertilizers |
| Dry grasslands | Grasslands with a low water budget (often on southern slopes) and a low nutrient level |
| Xerothermophile grasslands | Category of grasslands very dry and a with a poor nutrient level |
| Wetlands | Grasslands or pastures with a high water budget, wet most part of the year |
| Tall herbs | High vegetation, transition between forest and wetland |
| Bogs | Wetlands with accumulation of acid peat |
| Schrubs | Schrubs areas, like for the green alder (*Alnus viridis*) |
| Forest edges | Limits of forests |
| Forest | Any forest areas |
| Crops | Cultivated meadows |
| Garden and nursery | Anthropomorphic green areas |
| Anthropomorphic environments | Like roads borders, railways tracks, gravel pit |
| Fallen rocks | Any Fallen rock or boulder areas |
| River banks | Any river banks |

**Table S2:**
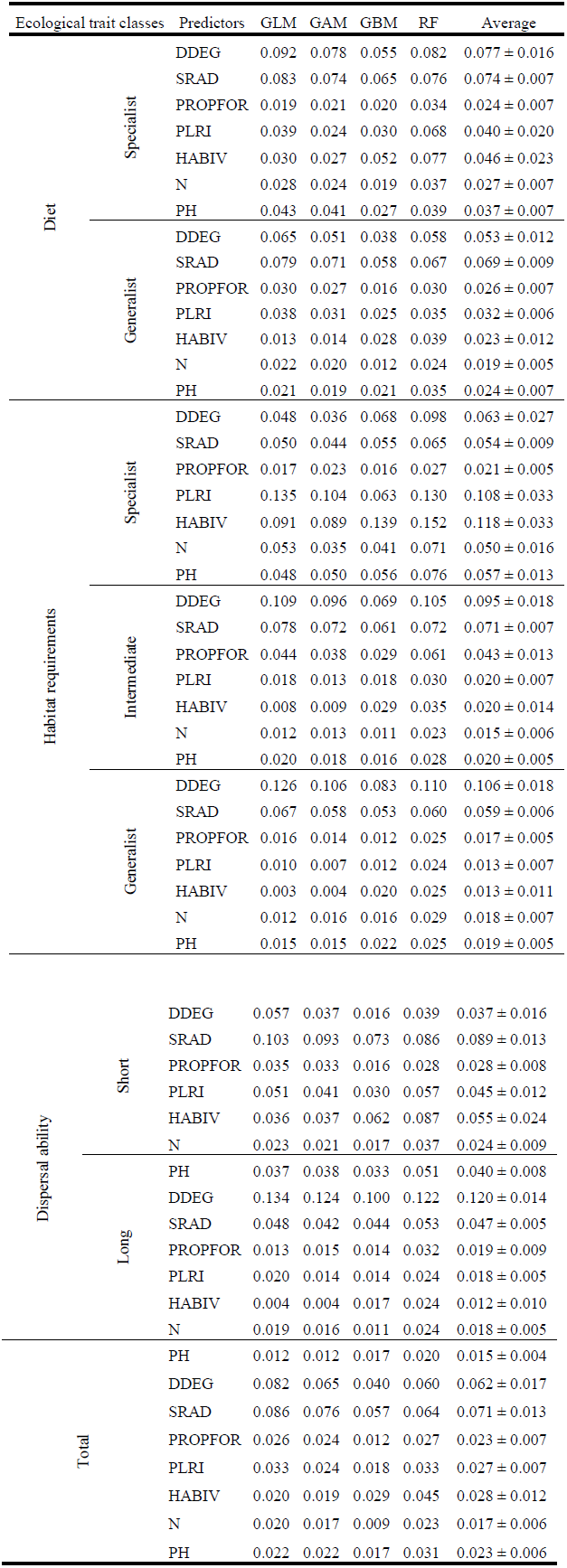
Results of the environmental predictor influence on butterfly species richness for the four statistical approaches investigated and following the ecological trait classifications.

**Table S3:**
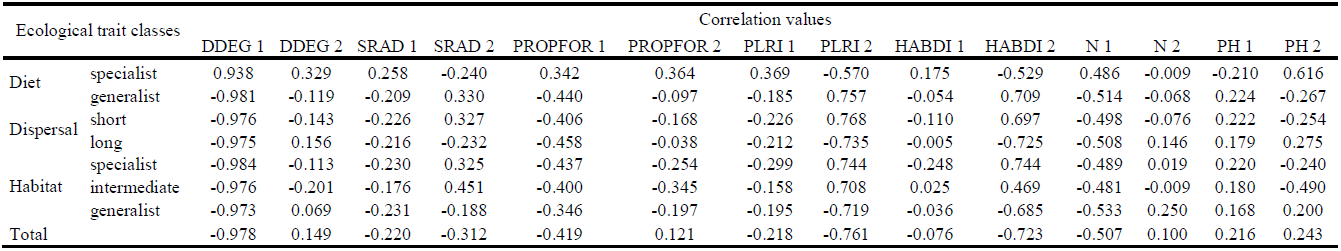
Correlation of the predictors with the first and second axis of the canonical correspondence analysis of butterfly species composition and according to the ecological traits investigated.

**Table S4:** Eigen values resulting from the canonical correspondence analysis of butterfly species composition for the seven different axes and according to the ecological traits investigated.

| Ecological trait classes |  | Eigen values |  |  |  |  |  |  |
| --- | --- | --- | --- | --- | --- | --- | --- | --- |
|  |  | axis<br>1 | axis<br>2 | axis<br>3 | axis<br>4 | axis<br>5 | axis<br>6 | axis<br>7 |
| Diet | specialist | 0.364 | 0.174 | 0.166 | 0.097 | 0.076 | 0.075 | 0.049 |
|  | generalist | 0.623 | 0.158 | 0.074 | 0.060 | 0.033 | 0.029 | 0.023 |
| Dispersal | short | 0.532 | 0.167 | 0.107 | 0.078 | 0.046 | 0.041 | 0.029 |
|  | long | 0.506 | 0.160 | 0.103 | 0.087 | 0.062 | 0.051 | 0.030 |
| Habitat | specialist | 0.462 | 0.182 | 0.115 | 0.079 | 0.061 | 0.056 | 0.044 |
|  | intermediate | 0.583 | 0.134 | 0.093 | 0.088 | 0.044 | 0.033 | 0.026 |
|  | generalist | 0.547 | 0.200 | 0.087 | 0.074 | 0.041 | 0.027 | 0.025 |
| Total |  | 0.525 | 0.165 | 0.102 | 0.079 | 0.049 | 0.043 | 0.038 |

